# Computed atlas of the human GPCR-G protein signaling complexes

**DOI:** 10.64898/2026.03.07.710286

**Authors:** Miglionico Pasquale, Matic Marin, Franchini Luca, Hiroki Arai, Nemati Fard Lorenzo Amir, Arora Chakit, Magda Gherghinescu, Natalia De Oliveira Rosa, Kise Ryoji, J. Silvio Gutkind, Cesare Orlandi, Asuka Inoue, Raimondi Francesco

## Abstract

Experimental mapping of G protein-coupled receptors (GPCR)–G protein signaling coupling has illuminated hundreds of receptors, yet the coupling specificity of a large fraction of this large receptor family remains unknown, thereby preventing the development of new GPCR-targeting therapies. Here, we used AlphaFold3 (AF3) to predict the 3D structures of the human GPCRome in complex with heterotrimeric G proteins. We used experimental GPCR-G protein binding data to show that AF3 predictions significantly discriminate between positive and negative binders, and used 3D structural features to train a machine learning (ML) algorithm to predict coupling potency. Interpretation of the ML model helped discriminate universal features governing the strength of G protein coupling from those determining binding specificity.

We computationally illuminated the coupling preferences of 180 non-olfactory GPCRs (non-OR) with previously unreported transduction mechanisms and experimentally validate the predicted couplings for multiple previously uncharacterized GPCRs, including QRFPR, GPR50, GPR37, GPR37L1 and GPRC5A. Our predictions established that Gi/o is the most prevalent coupling among non-OR GPCRs, which is often co-occurring with Gq/11 and, to a lesser extent, G12/13 signaling. Gs coupling is less common and restricted to specific clusters within the non-OR GPCRome phylogeny, likely due to stricter structural requirements for its binding. We also computed G protein complexes for over 400 ORs, establishing Gs as the most prevalent coupling. ORs are predicted to bind to Gs with a simpler interface compared to non-ORs, ultimately leading to energetically less stable complexes. Additionally, we predict recurrent bindings to Gq/11 and Gi/o proteins for ORs, suggesting potentially novel ORs signaling mechanisms.

We exploited the GPCRome coupling atlas to interpret healthy and cancer expression data, revealing the coupling of most GPCR-G protein co-expressed pairs. This analysis highlights a richer coupling repertoire in healthy tissues compared to cancer, likely reflecting the high signaling requirements of specialized normal cell functions, which are lost in most cancer cells due to their de-differentiated state or under cancer selection processes.

In summary, this study provides the first computational 3D atlas of the human GPCR-G protein transductome, thereby illuminating the signaling mechanisms of neglected GPCR classes and providing the basis for interpreting omics datasets from a myriad of pathological conditions, thus enabling the development of novel precision therapeutics.

## Introduction

The coupling of G protein–coupled receptors (GPCRs) to G proteins is one of the core mechanisms by which eukaryotic cells sense and respond to their environment and act as a workhorse for the evolution of multicellularity^1^. In humans, GPCRs represent the largest family of cell-surface receptors and regulate countless physiological processes, from neurotransmission and hormone signaling to immune response^2^. Their ability to integrate the signaling of a multitude of extracellular stimuli by selectively engaging distinct intracellular G proteins determines the specificity, magnitude, and outcome of cellular signaling. By elucidating how GPCRs recognize and activate distinct G protein subtypes in a given cellular context, it is possible to better predict signaling outcomes, explain drug effects, and design more precise therapeutics with fewer side effects^3,4^.

The human genome encodes for 16 α subunits of heterotrimeric G proteins, which are grouped into four major families, namely Gs and Gi/o, respectively activating and inhibiting Adenylate Cyclase signaling, Gq/11 activating phospholipase C and G12/13 activating Rho GTPases^5^. Qualitative information about GPCR transduction mechanisms is continuously reviewed and updated by the Guide to Pharmacology database^6^. In the last decade, many experimental protocols have been developed to quantitatively elucidate the G protein-coupling repertoire of GPCRs^7–15^. This interaction information is integrated and freely provided by databases such as GPCRdb^16^ and GproteinDb^17^, and it has been used to train machine learning (ML) predictors for GPCR-G protein selectivity, such as PRECOG^18^ and PRECOGx^19^.

Experimental structural determination is rapidly advancing to illuminate the structural basis of GPCR-G protein selective binding, with 1011 experimental structures released (as of September 2025) covering 286 unique GPCR-G protein pairs and 209 unique receptors. The comparative analysis of these structures highlighted the key roles of transmembrane (TM) TM5 and TM6 regions as selectivity filters^20–23^ and macro-switches for Gs vs Gi/o binding^24^, leading to alternative docking modes and different energetic stability^25,26^. The advent of AlphaFold (AF)^27^ has revolutionized structural biology, offering the possibility to predict the structure of proteins and their complexes from the linear amino acid sequences with near-experimental accuracy. We and others have leveraged AF-multimer to predict the 3D complexes of GPCR-G protein experimentally known to interact^17,26^, thereby enhancing the understanding of the structural basis for molecular recognition in the nucleotide-free complex. In particular, these studies allowed for a better understanding of the structural basis of GPCR-G protein recognition for less represented families, such as Gq/11 and G12/13 complexes^26^.

In this study, we build on our previous workflow and use AF3 to predict GPCR-G protein complexes across the entire human GPCRome, extending predictions to hundreds of receptors for which experimental information is still lacking, including the large family of Olfactory Receptors (ORs). To this end, we demonstrate, using quantitative GPCR-G protein datasets, that AF3 predictions can reliably discriminate positive from negative binders and can be used to predict reliable interactions at the complete human GPCRome level.

## Results

### AlphaFold 3 predictions discriminate between positive and negative GPCR-G protein coupling

We used the common coupling map from GproteinDb^17^ to establish a ground truth for GPCR-G protein interactions. This curated dataset aggregates binding data from multiple experimental sources, identifying 1,714 tested GPCR-G protein pairs, which involved 213 (54% of non-OR GPCRs) unique GPCRs and 13 G proteins α subunits. The dataset comprised 844 pairs (49%) classified as non-couplers (or negative binders, i.e., with normalized Emax equal to 0) and 870 pairs (51%) showing evidence of coupling (or positive binders, i.e., with normalized Emax > 0) (Figure 1A; Supplementary Table 1).

**Figure 1:**
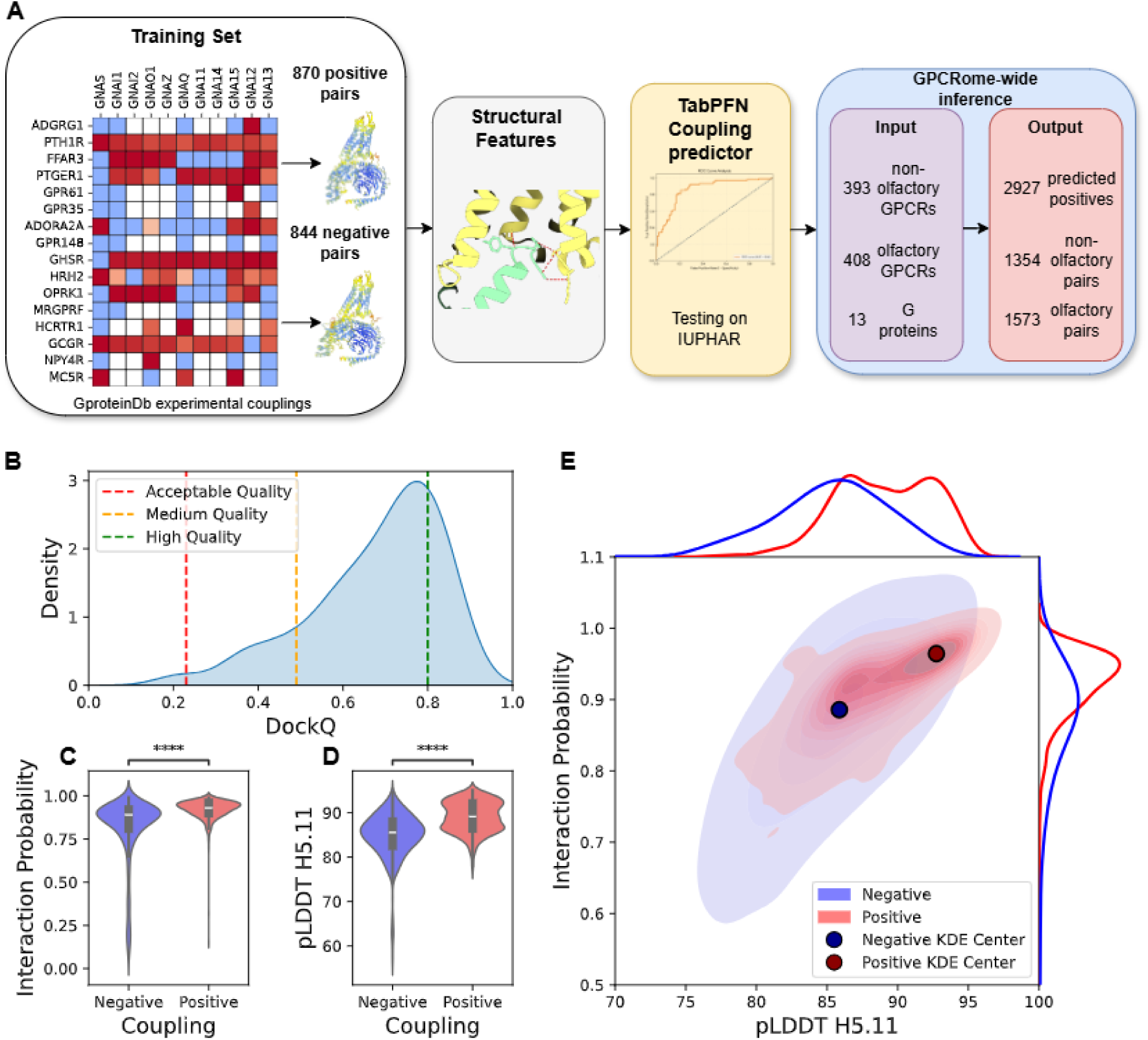
Computing GPCR-G protein couplings of non-olfactory GPCRome. A) Workflow of the GPCRome-wide coupling prediction. B) KDE plot of the DockQ of the AlphaFold 3 predictions with respect to the experimental structures released after the training cutoff. Violin plots of the Interaction probability (C) and the pLDDT of position H5.11 on the G protein (D) of positive and negative pairs. Boxplots show the median as the centre and first and third quartiles as the bounds of the box; the whiskers extend to the last data point within 1.5 times the interquartile range (IQR) from the box’s boundaries. The p-values have been computed with a two-sided Mann–Whitney U test (**** P < 0.0001); E) bi-dimensional KDE plot of the Interaction probability and the pLDDT of position H5.11 of positive and negative pairs.

To investigate the structural basis of these interactions, we predicted the structures of all 5,109 possible non-OR GPCR-heterotrimeric G protein complexes, considering each individual α subunit in complex with GNB1 and GNG1 subunits, using AlphaFold 3^28^ (AF3, see Methods). The vast majority of these predictions, 98.4% (5,032), were structurally plausible, correctly positioning the G protein at the intracellular, G protein-coupling interface of the receptor. The remaining 77 structures showed the G protein unrealistically docked to the extracellular side. Out of 13 mis-docked pairs with experimental coupling data, 12 are confirmed non-couplers, with only one (i.e., *APLNR-GNA15*) reported as coupling. We also considered the 213 predicted complexes for which experimental structures became available after AF3’s training cutoff. The comparison with the predicted models yielded a median DockQ^29^ score of the interface between the GPCR and Gɑ subunits of 0.73, with 98.6% (210) of the structures achieving at least acceptable quality based on established structural metrics (i.e., DockQ > 0.23^30^) (Figure 1B; Supplementary Table 2).

We investigated whether metrics and structural descriptors derived from AlphaFold3’s predictions correlate with experimental coupling data from GproteinDb. We derived an interaction probability between the GPCR and Gα chains from the residue-residue contact probability matrix (see Methods). We found it to be significantly higher for positive than negative binders (p-value=5.6E-87) (Figure 1C), as well as to be correlated with coupling (Spearman ⍴ = 0.551, p-value=1.3E-136; ROC AUC = 0.771; Supplementary TabSyle 1). Similarly, the ipTM score was overall correlated with coupling (Spearman ⍴ = 0.440, p-value = 6.2E-82, ROC AUC = 0.706; Supplementary Table 3), particularly for class A GPCRs (Spearman ⍴ = 0.485, p-value = 7.5E-85, ROC AUC = 0.735, p < 0.001 computed with bootstrap; Supplementary Table 4), showing that the interaction interface of coupled pairs was modeled significantly better than the one of non-couplers. We refined the predicted structures using Rosetta^31,32^, whose energy score captures information derived from statistical and physical principles, and characterized the energetic stability of the complex interfaces (see Methods). We found that Rosetta’s binding energy is also significantly more negative for positive vs. negative binders (p-value=2.2E-43) (Supplementary Figure 3), and correlated with coupling (Spearman ⍴ = 0.334, p-value = 6.0E-46, ROC AUC = 0.655; Supplementary Table 3), suggesting that coupled pairs are energetically more stable. We also derived a set of local (i.e., residue-level) structural descriptors, such as quality of the prediction (i.e., predicted Local Distance Difference Test - pLDDT), as well as the probability of contact for residues at the GPCR-G protein interface^26^. We found that residue-specific descriptors could also be associated with either positive or negative binding. For instance, the pLDDT of residues such as the G protein’s H5.11 (Common Gα Numbering (CGN)^33^), a conserved residue stabilizing G protein’s α5 helix intramolecular contacts, significantly discriminates between positive and negative binders (Spearman ⍴ = 0.505, p-value = 1.3E-111, ROC AUC = 0.762; Figure 1D; Supplementary Table 3).

By simply combining global features, such as AF-derived interaction probability, with local ones, such as G.H5.11’s pLDDT, it was possible to separate the distributions of positive and negative binders (Figure 1E). Overall, these findings suggest that the structural features derived from AlphaFold’s predictions could inform whether a predicted complex is a positive GPCR-G protein interaction.

### Training a structure-based predictor of GPCR-G protein selectivity

Inspired by these findings, we trained a supervised model (i.e., Precog3D) to predict the GPCR-G protein coupling from its 3D complex. We used a total of 184 global as well as local structural features from AF3-predicted complexes, which we input to TabPFN^34^, a foundational model for supervised learning on tabular data. We trained a regression model on 1.7k experimental interactions from GPCRdb (see Methods). We identified 4 as the optimal predicted binding strength cutoff to discriminate positive from negative binders (Supplementary Figure 2A). For testing, we considered a set of 316 couplings (114 positives and 202 negatives) that refer to 79 unique receptors from IUPHAR/BPS Guide to PHARMACOLOGY^6^ (GTPDB), which were not present in the training set.

Despite being a general model for GPCR-G protein coupling, Precog3D showed improved performances with respect to the state of the art predictors for GPCR-G protein coupling specificity (i.e., PRECOGx^19^), both in 5-fold cross-validation on the training set (Supplementary Figure 2B,C; Supplementary Table 1), as well as in testing (Figure 2A; Supplementary Table 5), when stratifying the predictions based on G proteins. Overall, the model reliably predicts GPCR-G protein coupling (ROC AUC = 0.82, Supplementary Figure 2D). To check whether these performances were inflated by some receptor or G protein class performing above average, we also evaluated the model’s ability to predict G protein binding at the individual receptor level. To this end, we tested the predictive performance of each receptor that was withheld from training, using a leave-one-out procedure (see Methods). We achieved a median accuracy above 0.8 for Class A (0.82), B2 (0.83) and C (0.86), while slightly lower for B1 (0.64) (Figure 2B). We also trained a similar model by leaving out the full list of couplings mediated by a specific G protein, which were only used for testing (see Methods). In this way, we forced the model to predict couplings to a specific G protein by complementarily learning them from all the others. Also in this case, the model achieved good performances, with several coupling groups with an AUROC above 0.8, such as GNAL, GNAS, GNA11, GNAQ, GNA14, GNAI1, GNAI2 and GNAO1 (Figure 2C).

**Figure 2:**
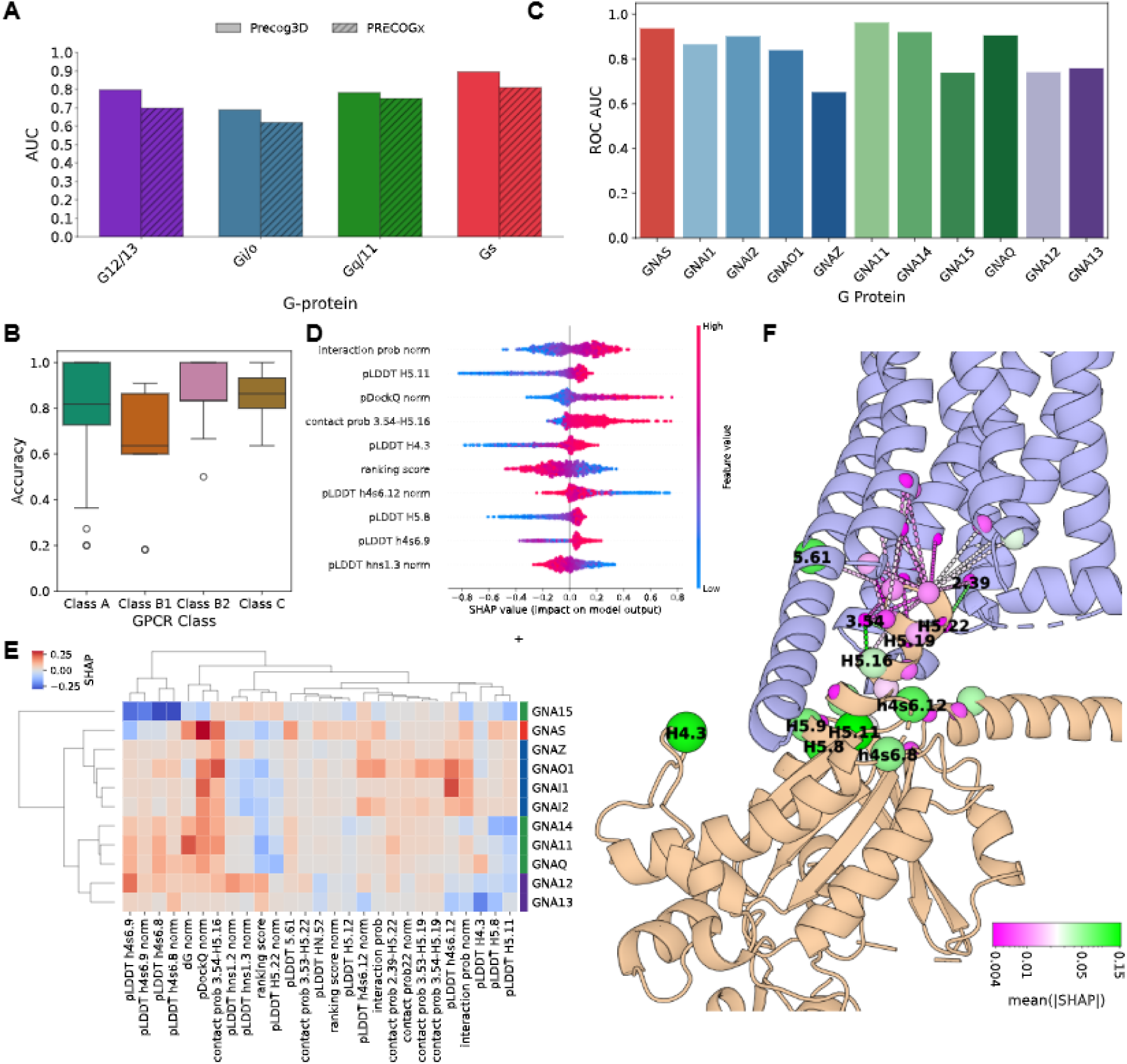
A machine learning model of Coupling specificity based on AF predictions. A) AUC ROC on the IUPHAR test set (G protein family wise); B) Receptor wise accuracy in the leave-one-GPCR-out setting. Boxplots show the median as the centre and first and third quartiles as the bounds of the box; the whiskers extend to the last data point within 1.5 times the interquartile range (IQR) from the box’s boundaries. C) ROC AUC in the Leave-one-G-protein-out setting; D) SHAP distribution of the 10 most important features (highest average absolute value of SHAP) in the IUPHAR test set; E) Clustermap of the average SHAP for each G protein in the positive pairs computed in an held-one-G-protein-out setting. Features with absolute value of the average SHAP bigger than 0.05 for at least one G protein are shown; F) 3D cartoon representations of a representative predicted structure (GPR21-GNAS) with the residues and the contacts highlighted if considered as local features. Sphere size and color indicate the average absolute SHAP on the testing set. Labels indicate the residues with mean(|SHAP|)>0.05.

We assessed the feature importance of the Precog3D model through a game theory approach, i.e., Shapley Additive exPlanations (SHAP)^35^ (see Methods). Interpretation of the general model revealed features that are universally important for predicting the intensity of G protein coupling (Figure 2D). In particular, the analysis confirmed the relevance of global features such as the Interaction Probability, Ranking score or normalized pDockQ score (Figure 2D). It also showed the importance of local structural features, from either the G protein (e.g., pLDDT H5.11, h4s6.12, H4.3, H5.8 and h4s6.9), the receptor (e.g., pLDDT 5.61, generic GPCR numbering^36^ or interface (contact probability 3.54 - H5.16) (Figure 2D,F). Analysis of the most important features in the model trained by leaving out for testing all couplings of a specific G protein revealed features that are more relevant to G protein coupling specificity prediction (Figure 2E). Indeed, unsupervised clustering on the average SHAP values obtained from positively predicted interactions within each coupling group recapitulated known phylogenetic relationships among the different G proteins (Figure 2E). In agreement with the explainability analysis of the general model, certain features, such as the pDockQ score or the contact probability of positions 3.54 - H5.16, universally and positively contribute to the binding prediction for most of the G proteins. Other features are specific to distinct G protein couplings. For instance, pLDDT h4s6.9 and h4s6.8 positively contribute to the binding prediction of GNA14, GNA11, GNAQ, GNA12 or GNA13, while they negatively contribute to the binding prediction of GNA15 and, to a lesser extent, GNAS. On the other hand, pLDDT h4s6.12, H5.8, H5.11, or contact probability 3.53 (3.54) - H5.19 positively contribute to Gi/o members (GNAI1, GNAI2, GNAI3, GNAO1, GNAZ) as well as to GNAS binding (Figure 2E). Overall, the most important positions for coupling specificity are found at the G protein’s H5, H4 and h4s6 loop, as well as at the receptor’s TM2, TM3, TM5 and TM6 intracellular extremities, spatially clustering around the G protein’s docking region (Figure 2F).

### Predicting GPCR-G protein coupling at the human proteome scale

We considered AF3-predicted 3D complexes of 180 human non-olfactory G protein-coupled receptors (non-OR GPCRs), as cataloged by GTPDB^6^, with 13 heterotrimeric G proteins, that were not yet considered in the training of Precog3D (see Methods). We then used the new model to discriminate predicted structures in coupled vs. not-coupled (Figure 1A). We mapped the predicted couplings onto a human GPCRome phylogeny that we constructed starting from AF-predicted, monomeric GPCR structures, by performing a structure-based MSA generation and phylogenetic analysis (i.e., FoldMason^37^, see Methods). The overall structural phylogenetics recapitulated known family relationships (Figure 3), assessed through the retention index^38^ (Supplementary Figure 3A), although, as expected, sequence-derived phylogenetics is more correlated with family classifications, likely due to historical reasons (Supplementary Figure 3A; Methods). We found no overall significant correlation between phylogenetics and coupling preferences, although the structural tree improves the correlation trend with respect to only sequence-based trees when considering experimentally known and predicted couplings for non-OR GPCRs (Supplementary Figure 3B). We found high consistency in coupling profiles and structural phylogenetics for certain families, such as Chemokine and Taste receptors, which are both predominantly bound to Gi/o (Figure 3). Other examples of coupling profiles correlating with structural phylogenetics are observed in Gs coupled receptors. For instance, the structural phylogram successfully segregates Trace amine-associated receptors, ꞵ-adrenergic receptors and other amine receptors (i.e., HTR4, HRH2, DRD1, DRD5, HTR6) that preferentially couple to Gs, from other amine receptors of the same families having preferences for Gi/o or Gq/11 (Figure 3). Several class B receptors comprising GLP1R and other secretin-like receptors are clustered together in the phylogram and show consistent coupling to Gs, as well as to other G proteins (Figure 3). Within the prostanoid and adenosine receptor families, the members that preferentially couple to Gs are clustered apart from members with different preferences (Figure 3). Other small clades grouping Gs-coupled receptors are the ones entailing Melanocyte-stimulating hormone receptors, as well as Class A orphans GPR3, GPR6, GPR12 and Bile receptor GPBAR1 (Figure 3).

**Figure 3:**
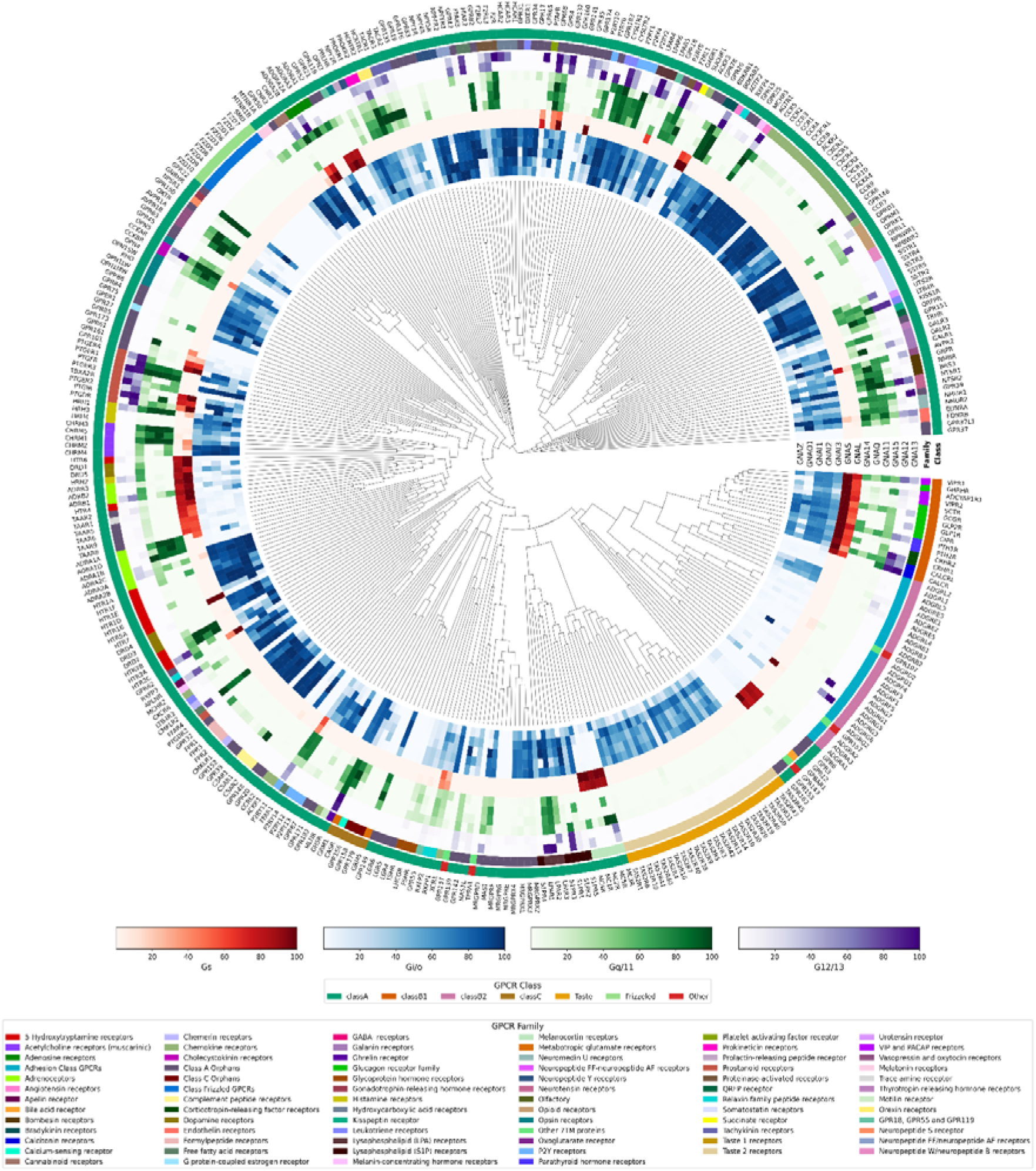
Structure-based phylogeny of the human GPCRome. The dendrogram displays receptor similarity obtained from structural alignments through FoldMason. Each receptor is annotated with the experimental (when available) or predicted coupling information for each individual G protein, in a color scale specific for the G protein family. Family and Class annotations from IUPHAR are also annotated through specific color schemes.

Overall, we found that predicted couplings for non-OR GPCRs recapitulate the trend of experimentally known binding^8,39^ (Supplementary Figure 4A), with Gi/o members being the most successful couplings, followed by Gq/11 ones (Figure 4A). Many receptors bind exclusively to Gi/o, which appears as the single, most frequent selective coupling, followed by combinations of Gi/o with Gq/11, Gi/o with Gq/11, G12/13, and Gi/o with Gq/11, G12/13, Gs. Selective couplings to other G proteins (e.g., Gs, Gq/11 or G12/13) seem to be a characteristic of a few receptors, both according to our model (Figure 4A) as well as experimental data (Supplementary Figure 4A).

**Figure 4:**
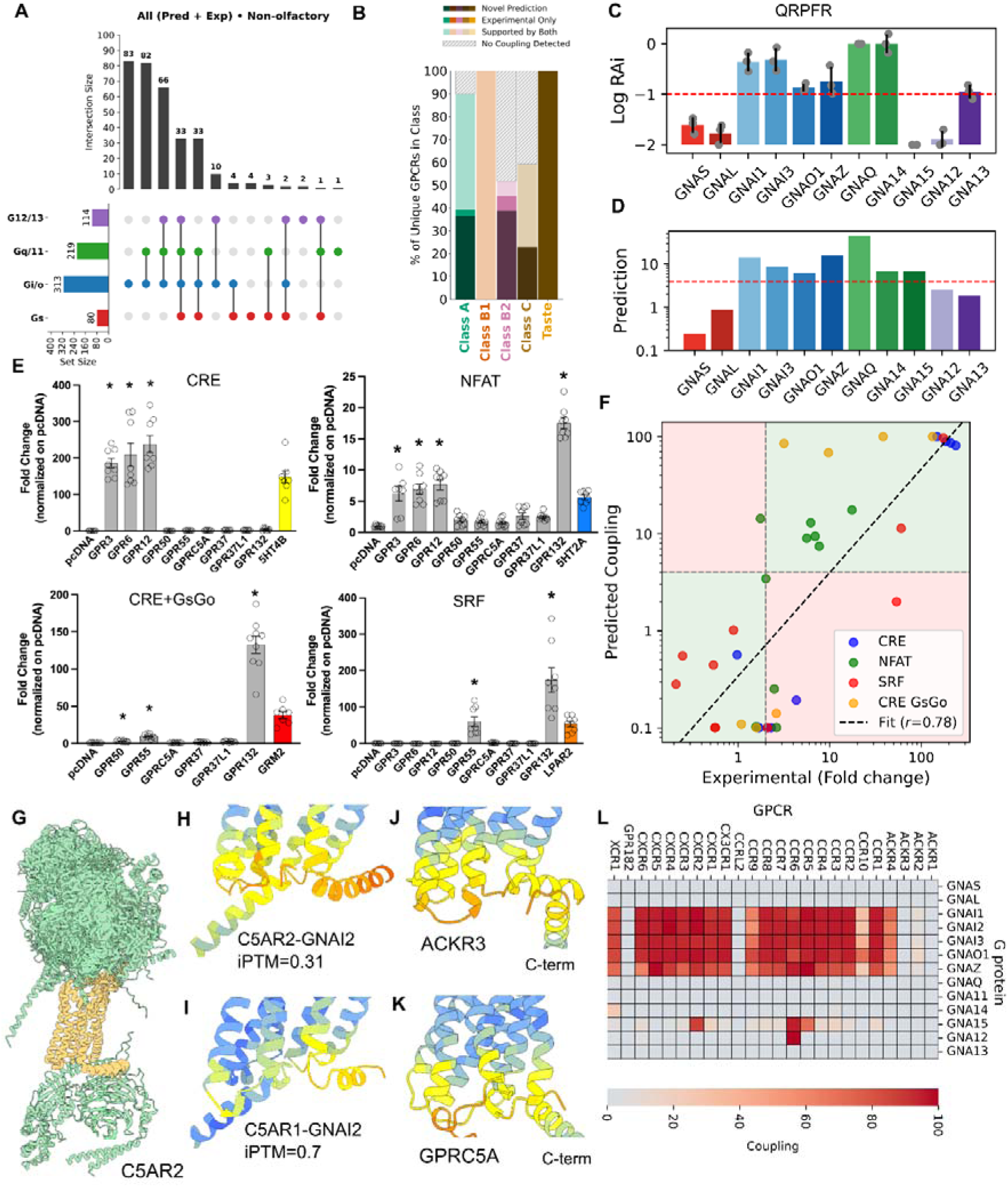
Inference at the non-OF human GPCRome and experimental validation. A) Distribution of the coupling combinations of the four G protein families in the non-OF human GPCRome (predicted and experimental); B) Fraction of GPCRs of each class with at least an experimental coupling among those present in the training set versus the fraction of GPCRs with at least a predicted or experimentally validated coupling in the entire GPCRome; C) G-protein-coupling profile of QRFPR assessed by the TGFα shedding assay. QRFPR was co-expressed with one of the Gαq chimeric constructs, in which the C-terminal sequence was substituted with that of the indicated Gα subunits, or with a C-terminally truncated Gαq construct (ΔC), together with the AP-TGFα reporter in the Gq/11/12/13-deficient HEK293 cells. LogRAi values were calculated from the concentration–response curves. Bars and error bars represent the mean and SD of three independent experiments; D) Precog3D predicted couplings for QRFPR; E) protein inducible luciferase reporters showing orphan GPCR constitutive activity, respectively for CRE, NFAT, CRE+GsGo and SRF. Data shown as fold change over control cells not expressing GPCRs. CRE, NFAT, CRE+GsGo were analyzed with Kruskal-Wallis vs pcDNA + Dunn’s multiple comparison while SRF was analyzed with one-way ANOVA + Dunnett’s multiple comparison (N=6-8 F). 5HT4B, GRM2, LPAR2, 5HT2A were used as positive controls for CRE, CRE+GsGo, SRF and NFAT reporters respectively, and were excluded from the statistical analysis; F) correlation between predicted coupling and luciferase reporter assay experimental data; G) Superimposition of the 13 predicted complexes of G proteins with C5AR2; H) Predicted complexes of C5AR2 and I) C5AR1 with GNAI2 (higher pLDDT in blue, lower in red); J) Predicted complexes of ACKR3 and K) GPRC5A with GNAI1; L) Heatmap of experimental and predicted couplings for chemokine receptors.

Predictions significantly expanded the number of transduction mechanisms across all non-OR GPCRs classes. A total of 87% Class A GPCRs have at least one predicted positive coupling (Figure 4B), which is comparable to the fraction of experimental bindings (86%), although predictions cover a larger number of receptors (overall 248 vs 151 experimental ones; Figure 4B). Class B1 (i.e., secretin-like GPCRs) and class C receptors have a slightly higher fraction of experimental vs. predicted bindings, mainly due to the larger pool of receptors considered for the predictions with respect to those experimentally assayed. Predictions for class B2 (i.e., adhesion GPCRs) doubled the fraction of receptors with at least one positive binding (0.6) compared to experimental ones (0.3) (Figure 4B). We also predicted positive couplings (i.e., mainly Gi/o proteins) for 100% of Taste receptors (Figure 3, Figure 4B).

Many predicted couplings are supported by literature evidence and Gene Ontology (GO)-terms annotations^40,41^. For instance, we found a total of 298 predicted interactions, lacking experimental evidence of binding, with a GO term matching the biological pathway corresponding to the bound G protein (“adenylate cyclase-activating G protein-coupled receptor signaling pathway” for Gs, “adenylate cyclase-inhibiting G protein-coupled receptor signaling pathway” for Gi/o, “phospholipase C-activating G protein-coupled receptor signaling pathway” for Gq/11 and “Rho-activating G protein-coupled receptor signaling pathway” for G12/13). The G protein interaction class with most couplings supported by GO is Gi/o (Supplementary Figure 4B).

We also predicted a total of 97 GPCR-G protein interactions whose 3D structures were experimentally determined after training of AF3 (Supplementary Figure 4B; Supplementary Table 6). Among these complexes, 82 (85%) are predicted to be positively coupled if we consider a predicted value greater than 1 (Supplementary Table 6). The discrepancy between structure and experimental binding data, however, is also present in the training set, since among the 117 training pairs with a newly resolved structure, 13 are annotated as negatively coupled. Nevertheless, for most of these complexes, we obtained high-quality predicted models across GPCR classes (Figure 1B).

We experimentally tested predicted couplings for several under-studied receptor systems. For QRFPR, AF3 predicted coupling to multiple members of Gi/o and Gq/11 families. The predictions are in line with transduction mechanisms through Gi/o and Gq/11 reported in GtoPDB, although quantitative coupling data are not available in GproteinDb (Figure 4C). Using the TGFα-shedding assay, we demonstrated a preference of QRFPR to couple with Gi/o members (GNAI1, GNAI3 and, to a lesser extent, GNAZ and GNAO1) as well as with Gq/11 members (GNAQ and GNA14) (Figure 4D, Supplementary Figure 5A; Supplementary Table 7). Our predictions showed good agreement with the TGFα-shedding assay, with a Pearson’s correlation of 0.66 (P-value = 0.026; Supplementary Figure 5B). We additionally tested predicted couplings for several orphan GPCRs (oGPCRs) using a luciferase reporter assay to evaluate the receptor constitutive activity (see Methods; Supplementary Table 8). We predicted that GPR50, an oGPCR with no experimental coupling information to date, would be a Gi/o selective receptor, which we confirmed experimentally (Figure 4E). We also predicted and experimentally validated couplings for oGPCRs whose signaling repertoire was only partially known. In detail, we predicted and showed experimentally that GPR3, GPR6 and GPR12 coupled to Gq/11, in addition to Gs (Figure 4E). The model also predicted Gi/o coupling for GPR3 and GPR6, which are consistent with IUPHAR data. Furthermore, we predicted novel, secondary couplings for receptors primarily known to couple to G12/13: for instance, we predicted and experimentally verified couplings of GPR55 to Gi/o, as well as of GPR132 to Gi/o and Gq/11 (Figure 4D; Supplementary Table 6,8). Overall, we found a Pearson’s correlation of 0.78 (P-value = 1.2E-8) between predicted coupling score and luciferase reporter assay log fold change (Figure 4F). Experimental validations also confirmed our forecasts for GPRC5A, GPR37 and GPR37L1, which we predicted not to couple to any G proteins (Figure 4E).

The model suggested multiple instances of atypical GPCRs, which seem widespread across the clades of the GPCRome phylogenetic tree, with a total of 87 receptors, across 14 families and 3 classes, predicted not to couple to any G protein (Supplementary Figure 4C; Supplementary Table 6). For instance, we predicted that C5AR2 (known to be a weak coupler^42,43^) is not coupled to any G protein. Indeed, for most of the complexes, the G protein is docked unrealistically on the receptor’s extracellular loops (Figure 4G). Only the complex between C5AR2 and GNAI1 is characterized by a plausible G protein docking topology, but with very low confidence, leading to a negative binding prediction (Figure 4H). In contrast, the close-paralogue C5AR1 is predicted to bind to several G proteins, including GNAI1, due to a more reliable interaction interface (Figure 4I). Predictions confirmed no G protein coupling for Atypical Chemokine Receptors (ACKRs) 1 and 3 (ACKR1 and ACKR3, Figure 4J,L; Supplementary Table 6), all characterized by conformations of the intracellular loops not compatible with G protein binding (Figure 4J). ACKR2 and ACKR4 are predicted to have residual couplings for Gi/o proteins (Figure 4L), likely due to higher structural similarity to chemokine receptors, which places them together in the same clade of the structural phylogenetic tree (Figure 3). ACKR1 and ACKR3 are structurally more divergent, resulting in occupying the phylogenetic tree far from the chemokine receptor’s clade. In particular, ACKR1 is found in a distal clade together with CCRL2 and GPR182, two receptors that are also considered atypical and indeed not predicted to couple to any G protein (Figure 4L). We also predicted several orphan receptors from class A and class C to be incompetent for G protein binding. For example, GPRC5 family receptors (e.g., GPRC5A, Figure 4K), which we experimentally validated (Figure 4E), as well as other Class C receptors such as GPRC6A and GABBR1, are all predicted to be atypical (Supplementary Table 6).

### The predicted signaling landscape of Olfactory receptors

We have systematically predicted through AF3 the 3D complexes of 408 Olfactory Receptors (ORs) and 13 heterotrimeric G proteins and predicted their coupling activity, finding a total of 1573 predicted positive couplings for 388 unique ORs (Figure 5A). Overall, ORs are characterized by distinct coupling preferences and by a simpler repertoire of coupling combinations (Figure 5A) compared to non-OR GPCRs (Figure 4A). In particular, ORs are predicted to be preferentially coupled to the Gs family (Figure 5A), specifically to GNAL (or Golf) (Figure 5A, Supplementary Figure 6A). These predictions are in line with the established knowledge that ORs signal through Golf to activate Adenylate Cyclase^44^. However, we also found a widespread, although weaker, coupling preference for Gq/11 members (Figure 5A), e.g., GNA15 (Supplementary Figure 6A), and Gi/o members (Figure 5A), in particular GNAZ and GNAO1 (Supplementary Figure 6A).

**Figure 5:**
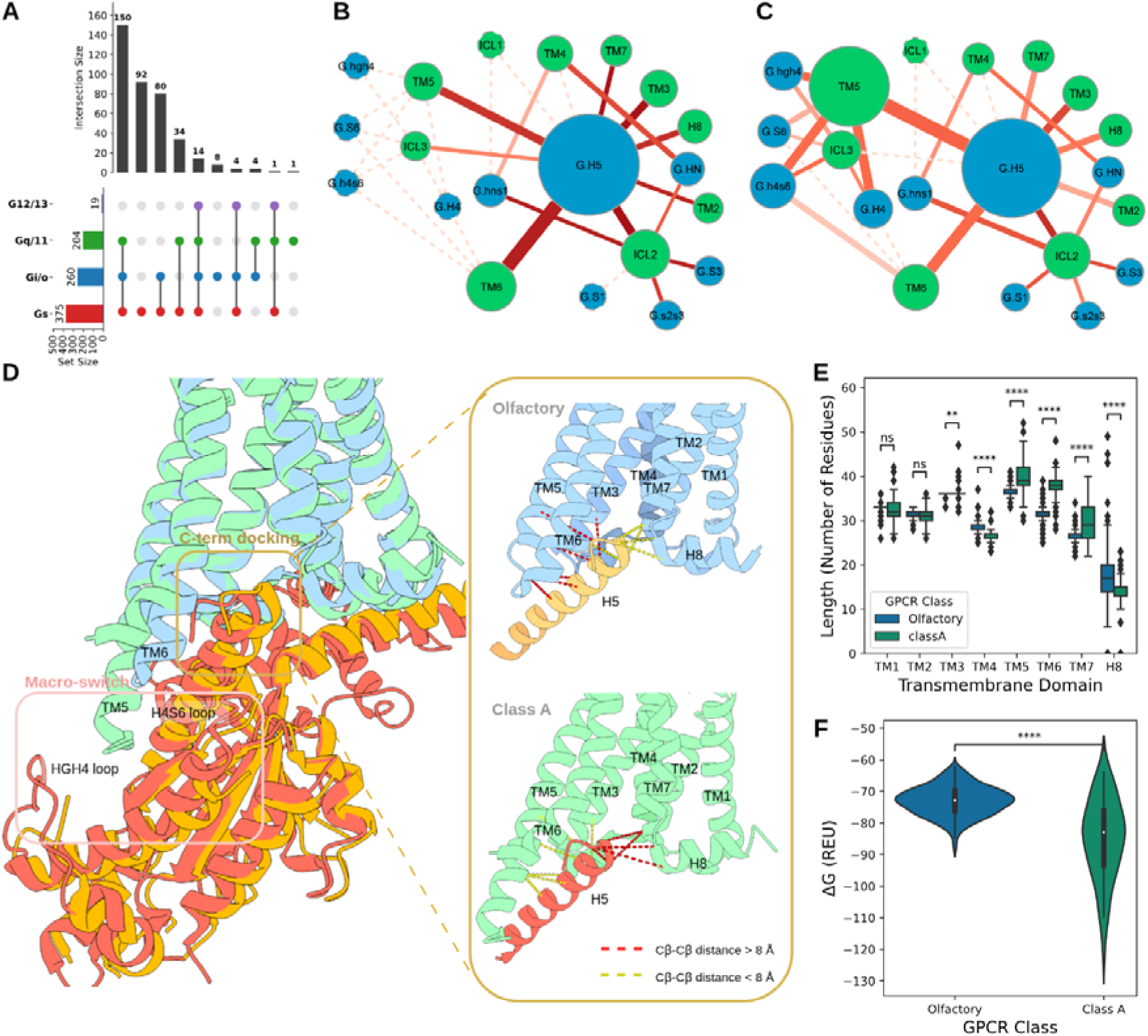
Prediction of the Olfactory receptors G protein couplings. A) Distribution of predicted coupling combinations of the four G protein families in olfactory receptors; B) SSE contact network for predicted Gs complexes for olfactory and C) class A GPCRs: GPCR and G protein nodes are colored in green and cyan, respectively. Node diameter is proportional to the total number of contacts mediated by that SSE. Edge thickness is proportional to the number of contacts between connected SSEs and coloring (darker red) is directly proportional to contact conservation; D) Superimposition of representative experimental structures of olfactory (PDB ID: 8F76) and class A (PDB ID: 3SN6) GPCRs in complex with Gs. Contacts present in only one of the two structures are highlighted in yellow when present (Cꞵ-Cꞵ distance < 8 Å) and in red when absent (Cꞵ-Cꞵ distance > 8 Å); E) Distribution of the length of the SSE in olfactory and classA receptors; F) Distribution of the ΔG binding (Rosetta Energy Units) in olfactory and class A receptors. Boxplots show the median as the centre and first and third quartiles as the bounds of the box; the whiskers extend to the last data point within 1.5 times the interquartile range (IQR) from the box’s boundaries. The p-values have been computed with a two-sided Mann–Whitney U test with Bonferroni correction (** P < 0.01, *** P < 0.001, **** P < 0.0001).

AF3 predictions revealed structural hallmarks of ORs binding to G proteins. We constructed a network of secondary structural elements (SSEs) derived from contacting residues at the 3D interface of GPCR-G protein complexes. By comparing the SSE networks of GNAS in complex with OR and non-OR GPCRs, we identified a major rewiring of contacts mediated by TM5 and TM6 with the G protein’s α5 helix (H5) (Figure 5B,C). In particular, TM6 makes more contacts with H5 (Figure 5B), and its importance is further remarked by the presence of a highly conserved motif (KAFSTCxSH-TM6.40) at the G protein’s interface, which is absent in non-ORs^45^. TM5 and ICL3 also contact H5, although to a lesser extent compared to non-OR GPCRs, while they lose multiple contact sites with hgh4, H4, S6 and h4s6 on the C-terminal lobe of the Ras domain, which are instead observed in the majority of non-ORs (Figure 5C). The predicted differences are consistent with available experimental structures (Figure 5D). Indeed, the superimposition of the only available complex between an OR (i.e., OR51E2) and a mini-Gs (PDB ID: 8F76) on the β2AR-Gs complex structure (PDB ID: 3SN6), shows that the G protein’s H5 helix is shifted towards TM2, TM7 and H8 in the OR complex, while it is closer to TM5 in the non-OR complex (Figure 5D). The centrality of TM5 in forming a second docking site for Gs in non-OR GPCRs is further corroborated by contacts with H4, as well as by the preceding and following loops, which are absent in the OR51E2-miniGs interface, due to partially unresolved hgh4 and h4s6 loops, which indeed confirms the absence of stabilizing contacts (Figure 5D). The alternative docking mode of Gs with ORs is connected to a significantly shorter TM5 (P-value = 1.5E-56), TM6 (P-value = 4.3E-90), and TM7 (P-value = 5.1E-5), as well as by significantly longer TM4 (P-value = 2.0E-94) and H8 (P-value = 6.4E-34; Figure 5E). These structural differences translate into different binding energetics, with ORs-Gs complexes that are energetically less stable than Class A paralogues (P-value = 7.5E-13; Figure 5F).

Overall, OR 3D models display a reduced interface diversity with Gs compared to non-ORs (Supplementary Figure 6B). The distinct predicted interfaces for ORs are further confirmed by unsupervised clustering of interface contacts, which separately cluster ORs and non-OR GPCRs when considering complexes with GNAL, GNAS or GNAI1 (Supplementary Figure 6C).

We extended the GPCRome phylogenetic tree by also considering OR structures (Supplementary Figure 7) and compared the structural phylogeny with the one obtained with standard sequence-based approaches. We found that the correlation of the coupling profile similarity with the distance from the structural phylogenetic tree (i.e., ⍴ = 0.49) is higher than the one obtained considering only non-OR GPCRs (i.e., ⍴ = 0.1), as well as the one obtained from sequence-based phylogram (⍴ = 0.22) (Supplementary Figure 3C). This result reinforces the notion that structures can provide more information about G protein coupling selectivity than sequence alone, and it also suggests that the coupling preference in ORs is more easily explained by sequence-structural relationships (Supplementary Figure 3C).

### GPCR-G protein coupling maps illuminate human tissue-specific signaling programs

We showcased the utility of the human GPCR-G protein coupling map to interpret transcriptomics datasets. To identify signaling axes that can transduce signals in a tissue-specific manner, we explored whether predicted GPCR-G protein pairs were also connected in co-expression networks obtained from vast bulk RNAseq datasets from healthy and cancer tissue samples (Figure 6A; see Methods). As an orthogonal validation, we found a high fraction of GPCR-G protein pairs co-expressed in healthy tissues, which are reported, or predicted, to couple by our structural model (Figure 6B). Indeed, out of 233 Class A GPCRs that are co-expressed with at least one G protein, 192 of them (82%) are also coupled (Figure 6B). Also, Class B1 GPCRs have a high fraction of co-expressed GPCR-G protein coupling pairs (88%; Figure 6B). A total of 28 ORs - G proteins are co-expressed in healthy tissues, out of which 27 (96%) are also predicted to positively couple (Figure 6B).

**Figure 6:**
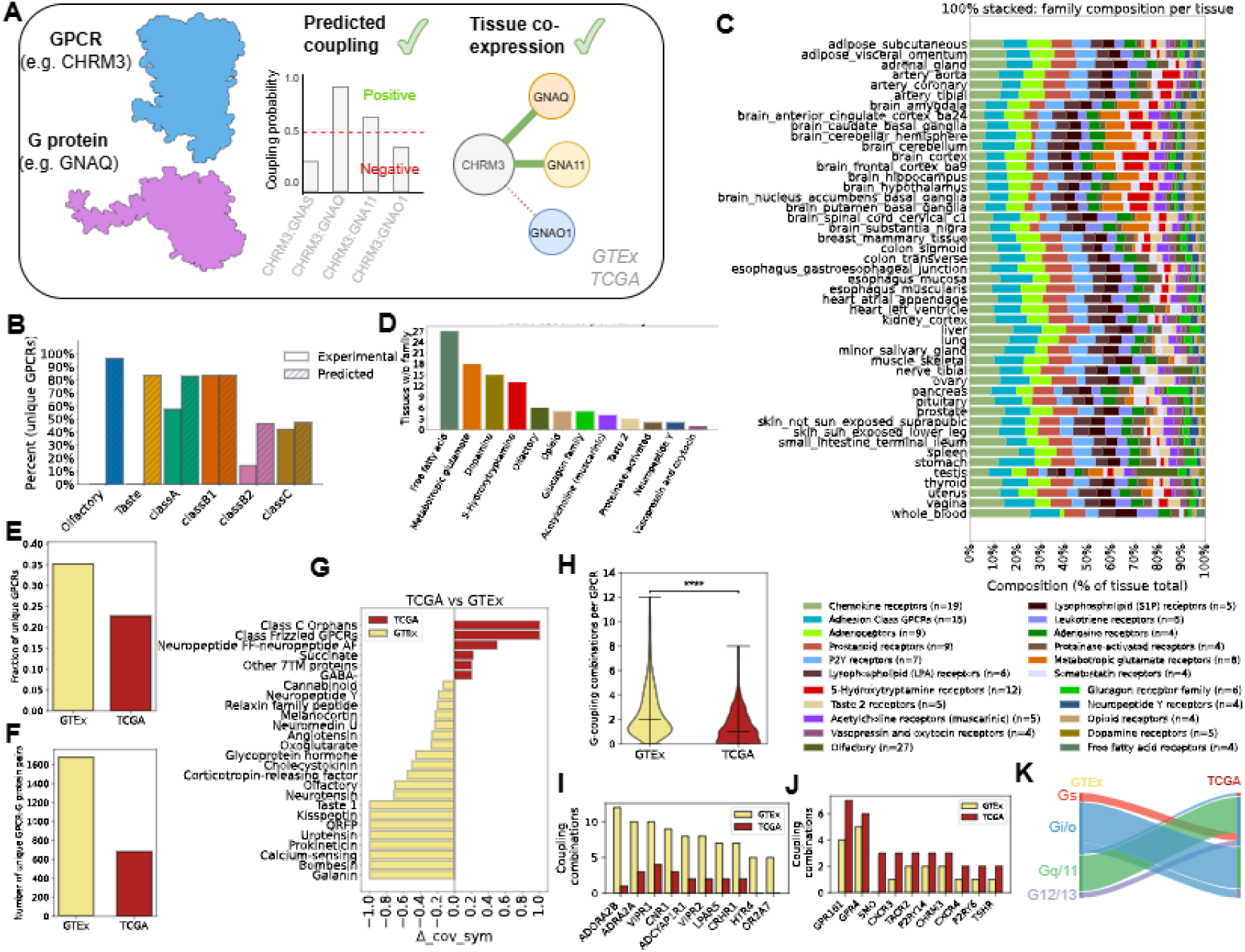
A) Human GPCRome-G protein transductome coupling and co-expression workflow B) Comparison of fraction of GPCR in different classes either predicted or experimentally validated to couple to G protein out of total GPCR in class reported (members that are co-expressed with (>0) and are coupling or predicted to couple to G protein(>4)) across all healthy tissue (GTEx) C) Comparison of fraction of GPCR families (members that are co-expressed with (>0) and are coupling or predicted to couple to G protein(>4)) in each healthy tissue (GTEx) D) Count of healthy tissues (GTEx) without specific GPCR families E) Fraction of unique GPCR (total=801) appearing across healthy tissues (GTEx) compared to cancer tissue (TCGA) (co-expression >0) F) Number of unique GPCR G protein coupled pairs appearing across healthy tissues (GTEx) compared to cancer tissue (TCGA) (co-expression >0); G) differential coupling of receptor families in TCGA vs GTEx assessed through the ΔSymm metric; H) Comparison of distributions of unique number of GPCR coupling combinations with different G proteins (**** P < 0.0001); I) Top 10 GPCRs based on the number of unique G protein coupling combinations in healthy tissue (GTEx) compared to cancer tissue (TCGA); J) Top 10 GPCRs based on the number of unique G protein coupling combinations in cancer tissue (TCGA) compared to healthy tissue (GTEx) K) Sankey diagram of rewiring of GPCRs coupling to four G protein families between healthy tissue (GTEx-left) and cancer tissue (TCGA-right).

Through G protein binding, it is possible to provide a comprehensive map of interacting GPCR-G protein pairs that are co-expressed in healthy tissues (Figure 6B or Supplementary Figure 8). Testis is the tissue with the highest number of co-expressed GPCR-G protein pairs, followed by lung and breast, while blood is the one with the smallest number of co-expressed interactions (Supplementary Figure 8A). Every tissue is characterized by a unique fingerprint of represented G protein couplings, although they follow the general distribution of G protein family coupling abundances, i.e., characterized by more Gi/o and Gq/11 couplings (Supplementary Figure 8A). Testis is also the tissue with the highest number of ORs-G protein bindings, followed by nerve tibial and spleen. While testis is characterized by OR couplings with Gs (GNAS), Gq/11 (GNA15) and Gi/o (GNAZ), spleen and nerve tibial are characterized by predicted OR couplings with members of Gs and Gi/o but not Gq/11 (Supplementary Figure 8B).

A higher degree of tissue-specificity emerges when considering the co-expression of all GPCR families with cognate G proteins. We found a more diverse repertoire of coupled GPCR families in healthy tissues (23 distinct families with at least four coupled members; Figure 6C) than in cancer ones (13 distinct families with at least four coupled members; Supplementary Figure 8C). In healthy tissues, families such as Free Fatty Acid receptors, Metabotropic glutamate receptors, Dopamine receptors, 5-Hydroxytryptamine receptors, and Olfactory receptors are characterized by the most tissue-specific co-expression patterns with cognate G proteins, i.e., quantified as the number of tissues missing such receptor families (Figure 6D). For instance, Metabotropic glutamate receptors are almost exclusively expressed in healthy brain tissues (Figure 6D). In cancer, in addition to the brain, they are also found co-expressed in tissues such as the pancreas, stomach and esophagus (Supplementary Figure 8C, Supplementary Table 9).

Overall, we found a higher number of GPCRs co-expressed with cognate G proteins in healthy tissues, both when considering unique receptor genes (Figure 6E) or GPCR-G protein pairs (Figure 6F). Certain receptor families are preferentially co-expressed in cancer or healthy tissues. Families such as Galanin, Bombesin and Calcium-sensing receptors are co-expressed with their coupled G proteins in several healthy tissues but not in cancer (Figure 6G). For instance, Galanin receptors are exclusively co-expressed in healthy brain (GALR1-Gi/o proteins), colon (GALR1-Gi/o and GALR2-Gq/11 or Gi/o proteins) tissues, but never in their cancer counterparts (Figure 6G; Supplementary Table 9). On the other hand, Class C Orphans, Frizzled GPCRs, Succinate receptor and Neuropeptide FF receptors are among the receptors that are co-expressed with G proteins in cancer but not in healthy tissues. Frizzled receptors, in particular SMO, are exclusively expressed in cancer tissues, particularly colon and lung (i.e., together with Gi/o and GNA12 proteins), while they are never found co-expressed in any healthy tissue (Figure 6G; Supplementary Table 9).

The greater diversity of GPCR coupling maps in healthy tissues compared with cancer tissues is also evident from the number of G protein family combinations, co-expressed with their cognate receptors (i.e., *coupling repertoire*). Indeed, in healthy tissues, we observed a significantly higher coupling repertoire than in cancer, considering GPCRs and tissues altogether (P-value = 7.6E-20; Figure 6H). For the vast majority of GPCRs, we observed a reduction of coupling repertoire in cancer tissues compared to healthy ones, with 201 out of 284 (70%) receptors showing a reduced repertoire across tissues compared to only 15 receptors (5%) showing an increased G protein repertoire complexity (Supplementary Table 9). In healthy tissues, a total of 162 receptors are characterized by more than one tissue-specific, coupling repertoire combination, while in cancer, such coupling promiscuity characterizes approximately half of the receptors (90) (Supplementary Table 9). For instance, ADORA2B is characterized by 12 distinct combinations of coupled G proteins that are co-expressed in 36 healthy tissues, while in cancer, it is co-expressed only with Gq/11 in 19 unique tissues (Figure 6I; Supplementary Table 9). On the other hand, SMO and GPR161 are among the receptors with the largest increase in the repertoire of co-expressed G proteins in cancer compared to healthy tissues (Figure 6J).

This indicates that the coupling repertoire in healthy tissues is highly diverse, and that switching between different signaling configurations, for the many promiscuous receptors, seems to be a widespread, common phenomenon in health, allowing extracellular signals to be contextualized to tissue-specific patterns of co-expressed G proteins. This diversity is greatly lost in cancer, with fewer examples of receptors gaining coupling repertoire relative to their healthy tissue counterparts.

The switching among alternative coupling configurations for promiscuous GPCRs also happens when comparing cancer with healthy tissues. Indeed, many GPCRs co-expressed with different cognate G proteins switch their predicted configuration towards Gi/o proteins in cancer. On the other hand, many Gi/o-coupled GPCRs in healthy tissues switch to Gq/11, followed by G12/13 and Gs (Figure 6K). In particular, we found many instances of Chemokine and P2Y receptors switching co-expressed, coupled G proteins from GNA15 in healthy tissues to GNAI2 in multiple cancers (Supplementary Figure 9).

## Discussion

AI has revolutionized protein structure prediction, reducing the gap between structural biology determinations and genome sequencing, enabling the structural description and visualization of the vast repertoire of biomolecules that are profiled at the genomics scale. In this study, we have used AF3, the state-of-the-art algorithm for protein structure prediction, to forecast the 3D structures of the human GPCRome in complex with heterotrimeric G proteins, thereby providing the first high-confidence structural map of GPCR transduction mechanisms, which we used to illuminate large-scale transcriptomics datasets.

Although AF3 has shown very good performance in predicting biomolecular assemblies, predicting protein complexes remains an open problem due to the huge search space of multimeric complex interfaces^46^. Specifically for GPCRs, ligand-receptor docking through AF is still an open problem^47^, due to the large variability not yet fully explored in experimental structures, with both successes^48^ and failures^49^ reported. On the intracellular side, we have previously shown that AF reliably predicts 3D GPCR-G protein complexes^26^, possibly due to a more conserved interface topology that has been accurately learnt by the AF model. Indeed, in the present study, we further validated the quality of the predictions against experimental structures solved after AF3 training, showing that the vast majority of them (98% out of 219 structures) are predicted with a quality DockQ score greater than the established threshold of 0.23 (Figure 1B).

Another open question posed by AF-multimer predictions is whether the quality of the structural predictions can discriminate between positive and negative direct interactions. This task is hindered by the lack of true negative binder information affecting most interactomics datasets^50^. Earlier studies showed that AF can discriminate between interacting and non-interacting proteins with high specificity^51^. Recently, a deep-learning framework that incorporates co-evolutionary information from deep multiple sequence alignment in combination with AF2 has shown good precision and recall performances in discriminating high-confidence PPI positive interactions from randomly-defined negative pairs with no evidence of interaction^52^. Another recent method, called SPOC, used AF predictions of a curated set of both positive and negative interactions to train an ML model to identify high-confidence PPIs^53,54^. Similarly, an ML method has been recently developed, based on AF predictions, to predict E2–E3 ligase binding specificity^53,54^.

In the GPCR field, multiple datasets have been generated by systematically profiling thousands of GPCRs-G protein interactions^8–15^, providing a wealth of binding information that can be used to accurately identify both positive and negative binders, which represents a key quantitative information mostly missing from the training sets of previous efforts to predict PPI based on AF models. We have systematically predicted 3D binary complexes for both positive and negative receptor-G protein couplings from these datasets, and correlated AF-derived prediction scores and structural features with coupling activity, showing that they can statistically discriminate interactions. Indeed, the new model (Precog3D), which we trained on AF3 models-derived structural features, accurately predicted GPCR-G protein coupling activity and showed competitive performances in predicting G protein binding specificity when compared with PrecogX^19^. The general improvement of performance is remarkable, if we consider that Precog3D was trained on the entire set of experimental interactions to predict binding strength, whereas Precog(X) consisted of separate models trained on G protein-specific binding data. The model also showed good G protein-coupling discrimination when trained by leaving out either entire GPCR- or G protein-specific couplings, further confirming the robustness of predictions based on AF3 complexes. The predictions for certain G proteins, such as Gi/o members, still pose some difficulties, likely due to the ease of binding for this protein family, which makes the training and testing sets “positive prone”.

Our predictions correlated well with experimental TGFα shedding assay data^7,8^ (Spearman ⍴ = 0.514, p-value = 1.9 E-48; Supplementary Figure 3), which have been largely removed from GproteinDb with the only exception of GNAQ and therefore can be regarded as an additional validation set. This value is comparable to the correlation between GproteinDb and the TGFα shedding assay itself, likely reflecting the inherent noise and batch effects in experimental coupling measurements (Supplementary Figure 3). Such inconsistency across different experimental binding assays is also observed in experimental structural determinations, with over 20 experimental structures corresponding to interactions in the training set that were reported as negatives in GproteinDb (Supplementary Table 1,2). Despite these biases, the model was able to correctly predict as positives 82 out of 97 (85%) experimental structures that were determined after training the AF3 model. Feature importance analysis of the general model highlighted structural characteristics that are universally important for G protein binding strength, either globally, such as binding probability, or locally, such as the probability of specific residues, such as G protein position H5.11. The importance identified by the coupling models of conserved positions such as H5.8 and H5.11, which lie on the α5 helix’s “transmission module”^33^, is likely connected to their role in mediating intra-molecular contacts important for transmitting the receptor’s activation signal to the G protein’s nucleotide binding pocket. Indeed, contact analysis of these positions in available GPCR-G protein complexes revealed G protein-specific signatures (Supplementary Figure 2E), which are captured by our model. Feature importance in the leave-G protein-out models also highlighted features associated with coupling specificity, which confirmed several positions previously identified based on contact analysis of experimental structures (e.g., 3.53 (3.54) - H5.19)^24,26^.

Our resource enabled us to expand the repertoire of G protein-coupling at the human GPCRome scale. We shed light on the coupling preferences of many orphan receptors, including several Class A and Class C orphan receptors, as well as the entire olfactory receptor family. The structure-based prediction model also explained the basis of signaling for atypical receptors, e.g., ACKRs, and suggested new receptor instances, similarly predicted not to bind G proteins, across families and classes. Importantly, predictions for the entire non-OR GPCRs group confirmed the experimentally observed tendency to couple with Gi/o, either individually or in combination with other G proteins, particularly Gq/11, G12/13 and, to a lesser extent, Gs. Exclusive, non-promiscuous coupling to non-Gi/o proteins in non-OR GPCRs appears as a rare evolutionary event (Figure 3, 4A). Moreover, Gs coupled receptors are characterized by structural features (e.g., TM5-TM6 selectivity filter) that lead to their clustering in specific clades of the GPCRome phylogeny, either intra- or inter-family.

Instead, ORs have a strong predicted tendency to couple with Gs proteins (particularly GNAL). Inspection of the structural models revealed that OR complexes with Gs substantially differ from non-OR GPCRs at the level of the lengths of the TM5, ICL3, TM6, TM7 and H8, and are in turn characterized by less energetically stable complexes. We speculate that a transient, weaker binding of ORs to Gs might be correlated with the different odorant binding mode, which, particularly for OR class II, is mainly characterized by Van Der Waals interactions and fewer H-bonds or salt-bridges^55^. The weaker G protein binding might contribute to eliciting a more nuanced response to each odorant, which has to be combined and integrated with a multitude of different odorant stimuli, perceived by different receptors and neurons. We also speculate that such structural differences are connected to the widely different G protein coupling repertoire for ORs compared to non-OR GPCRs. Indeed, although simpler and centered on Gs, ORs’ repertoire also entails combinations with Gi/o and Gq/11, which are predicted, although to a lower strength, for many ORs. Although unexpected, several of these non-canonical OR signaling have been reported in the literature. For instance, GNAO1 has recently been proposed to mediate alternative, inhibitory pathways in olfactory neurons^56,57^. Gαo has also been reported as the primary G-protein α-subunit mediating the detection of peptide and protein pheromones by sensory neurons of the vomeronasal organ^58^. We expect that our emerging coupling predictions will help propel further research on the repertoire of signaling mechanisms elicited by the large OR family.

Although we found no strong correlations between GPCR phylogenetic trees and coupling preferences, we demonstrated that GPCR similarities obtained from structural alignment of the receptors improve correlations with couplings compared to sequence-based methods only, as well as they might provide new opportunities to improve the classification for orphan GPCRs. This is particularly evident for certain families or subfamilies, such as Taste and Chemokine receptors, whose cladogram is highly consistent with Gi/o coupling preferences, as well as amine receptors, class B receptors, and ORs specific for Gs couplings. The close correlation observed between the structural phylogeny and coupling specificity of certain families is likely linked to evolutionary trajectories constrained to specific cellular and tissue contexts (e.g. Taste or Odorant receptors).

In the current study, we have modeled only the binary complex between receptor and heterotrimeric G proteins, considering the methodological limitations in reliably docking ligands, and the fact that for orphan GPCRs the information about their ligands is missing. In the future, it will certainly be interesting to assess the capability of AF, as well as of our model, to predict G protein coupling bias by also modeling the binding of the ligand as well as the conformational transition linked to the receptor’s activation. Another important biochemical information missing from the current model is the representation of the complexity of GPCR post-translational modifications^59–61^, such as phosphorylations, which might also affect the binding of G proteins and the recruitment of ꞵ-arrestins. Indeed, future efforts will also need to address additional intracellular transducers such as arrestins and receptor-modified states.

We employed the human GPCRome coupling atlas to interpret co-expression networks from bulk-RNAseq data from healthy and cancerous tissues. We found high fractions of GPCRs co-expressed with G proteins that are either experimentally reported or predicted to couple. This suggests that co-expression networks capture plausible biologically relevant regulatory interactions. Indeed, every tissue retains a highly specific pattern of co-expressed GPCR-G protein pairs, with certain families selectively expressed or enriched in specific tissues. Overall, we found a higher diversity of co-expressed receptor-G protein couplings in healthy tissues than in cancer. Taken together, this suggests that a richer repertoire of signaling competent GPCR-G protein axes is required in healthy tissues, likely reflecting the more specific signaling and cell-communication requirements of specialized cells and tissues. In cancer, such specialization is lost, likely due to the de-differentiated nature of cancer cells and selection pressure driving cancer evolution. Supporting the latter, certain GPCR-G protein axes are prevalently co-expressed in cancer. For instance, we found no co-expression of SMO in healthy tissues, consistent with its role in the early development of multiple organs^62,63^, whereas it is expressed in multiple cancers. Inhibition of SMO is one of the key targeted interventions to block the hedgehog signaling pathway in tumors such as medulloblastoma and basal cell carcinoma^64^. Similarly, GPR157, Succinate receptors (i.e., SUCNR1) and Neuropeptide FF-neuropeptide AF receptors (i.e., NPFFR1) are mainly co-expressed in cancer, thereby exposing novel candidate anti-tumor targets. Indeed, Succinate receptors have recently been highlighted as new drivers of cancer metastasis^65,66^, as well as new targets for cancer immunotherapy^67^. SMO, together with GPR161, another master regulator of the sonic hedgehog pathway, is one of the few GPCRs whose repertoire of coupled G proteins is increased in cancer tissues. At the opposite end, we found ADORA2B as the receptor with the greatest reduction in G protein-coupling repertoire between healthy and cancerous tissues. Intriguingly, agonistic treatment of ADORA2B has been recently proposed as an alternative strategy to counteract oncogenic SMO activity in basal cell carcinoma^68^.

We speculate that the combination, or sequential application of GPCR drugs, through the emerging notion of nudge, or state-shifting drugs^69^, could be leveraged to rewire GPCR signaling to recover the healthy G protein repertoire, therefore neutralizing the effect of the tumor microenvironment. The possibility to model ligand-biased GPCR-G protein signaling complexes, integrated with cellular-level omics datasets and cellular response to drug perturbations, will offer the possibility to finely modulate signaling networks to revert disease states through GPCR targeting.

## Methods

### Datasets

The list of human non-olfactory GPCRs was retrieved from the target and family list downloaded from IUPHAR/BPS Guide to PHARMACOLOGY (version 2025.02), selecting entries of type “gpcr” and excluding those annotated as “pseudogene” in the “HGNC name” field.

The list of human olfactory GPCRs was assembled by combining two sources. First, entries were retrieved from UniProt^70^ by filtering for proteins annotated with “Olfactory” AND “HUMAN”, excluding entries containing “pseudogene”, and requiring the Pfam domain PF13853 (olfactory receptor). In parallel, the human olfactory receptor list was obtained from GPCRdb^16^. The two lists were then merged and deduplicated to generate a final non-redundant set containing all unique receptors present in either source.

The selection of experimental structures was done as in ^26^. Briefly, we utilized the mapping between PDB^71^ and Pfam^72^ provided by SIFTS^73^ (2025/09/04 update) to retrieve all structures containing both a GPCR and a G protein α subunit. We used the Pfam entry PF00503 to identify structures of Gα subunits, and Pfam entries PF00001 (rhodopsin receptor family - class A), PF00002 (secretin receptor family - class B), PF00003 (class C receptors), PF01534 (Frizzled/Smoothened family) and PF13853 (Olfactory receptor) to identify GPCRs. We found 1011 structures that met this criterion. If more than one GPCR or Gα chain were in the structure, we considered the pair of chains with the highest number of contacts between them (Supplementary Table 1).

The Generic GPCR residue numbers^36^ were extracted from GPCRdb^16^.

### GPCR-G protein complexes prediction via AlphaFold 3

We used AlphaFold 3^28^ (v3.0.1) locally to generate 3D models of GPCR-heterotrimeric G protein complexes. The human canonical protein sequences of GNB1 and GNG1 were used to model the β and γ subunits, respectively. GPCR N-termini were truncated to a maximum of 50 residues upstream of TM1, and C-termini to 100 residues downstream of helix H8, to reduce terminal disorder. We considered only templates released before 30 September 2021 (the same cutoff used for the training of AlphaFold 3) and used 0 as the seed for all predictions. Among the 5 models generated for each GPCR-G protein complex, only the one with the best AlphaFold-generated ranking score was considered for further analysis.

We then employed AlphaFold’s confidence score (predicted Local Distance Difference test - pLDDT) to remove residues predicted with low confidence, which might lead to artefactual contacts. Hence, we removed all the residues with pLDDT <70 with the exception of the 7 C-terminal residues of the Gɑ subunit, which were always kept in the structure, since they are known to be important for specificity^8^.

Comparison of GPCR-G protein predicted models with experimental structures was performed using the pip-installable DockQ package^29^ (v2.1.3). For this comparison, we considered 583 structures deposited in PDB after 30 September 2021 and containing GPCR-G protein pairs without an experimental structure deposited before the same date. Only the DockQ of the interface between the receptor and the Gɑ subunit was considered. If more than one experimental structure of the same GPCR-G protein pair was found, only the one with the highest DockQ was considered.

### Interface analysis with Rosetta

To analyze the GPCR-Gα interface in a 3D structural model, we first relaxed the structure using the Rosetta relax application^31^ with the default fast relax options (5 cycles) without constraints, then we ran Rosetta InterfaceAnalyzer^32^ (https://www.rosettacommons.org/docs/latest/application_documentation/analysis/interface-analyzer), from the RosettaCommon software suite (version 3.14), specifying the chains of the GPCR and the Gα that are interacting in the complex. The interface energy and the ΔSASA are calculated as the difference in energy and Solvent Accessible Surface Area (SASA) in the bound and unbound structures. We ran InterfaceAnalyzer with the “-pack_input” and “-pack_separated” flags to optimize the side chain configuration before and after separating the chains.

### Features description

We considered a total of 92 features derived from the AlphaFold 3 structures as described in the following table (Table 1).

**Table 1:**
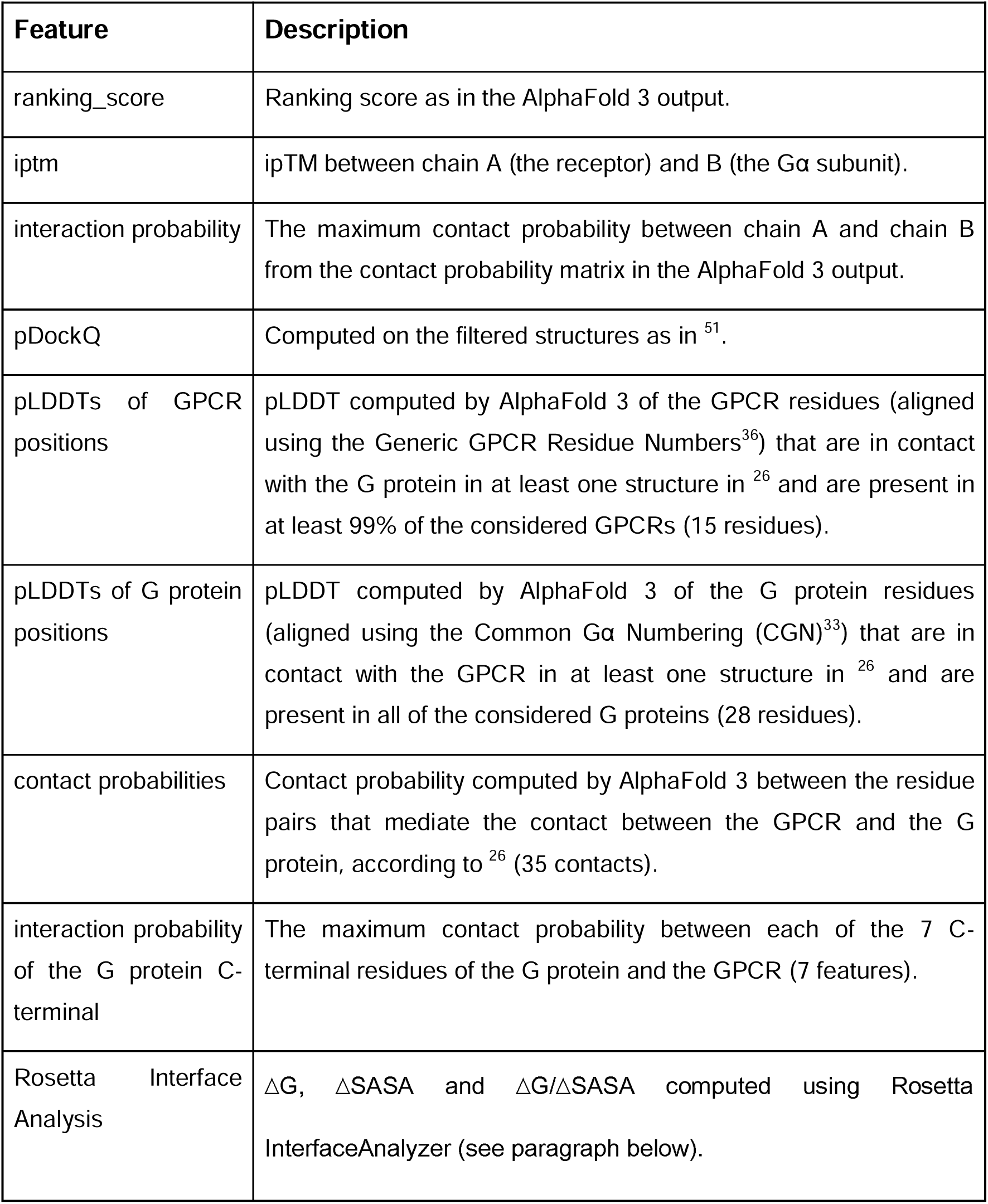
Structural features used for training the machine learning model.

| Feature | Description |
| --- | --- |
| ranking_score | Ranking score as in the AlphaFold 3 output. |
| iptm | ipTM between chain A (the receptor) and B (the Gα subunit). |
| interaction probability | The maximum contact probability between chain A and chain B from the contact probability matrix in the AlphaFold 3 output. |
| pDockQ | Computed on the filtered structures as in <sup>51</sup>. |
| pLDDTs of GPCR positions | pLDDT computed by AlphaFold 3 of the GPCR residues (aligned using the Generic GPCR Residue Numbers<sup>36</sup>) that are in contact with the G protein in at least one structure in <sup>26</sup> and are present in at least 99% of the considered GPCRs (15 residues). |
| pLDDTs of G protein positions | pLDDT computed by AlphaFold 3 of the G protein residues (aligned using the Common Gα Numbering (CGN)<sup>33</sup>) that are in contact with the GPCR in at least one structure in <sup>26</sup> and are present in all of the considered G proteins (28 residues). |
| contact probabilities | Contact probability computed by AlphaFold 3 between the residue pairs that mediate the contact between the GPCR and the G protein, according to <sup>26</sup> (35 contacts). |
| interaction probability of the G protein C-terminal | The maximum contact probability between each of the 7 C-terminal residues of the G protein and the GPCR (7 features). |
| Rosetta Interface Analysis | $\Delta G$, $\Delta SASA$ and $\Delta G/\Delta SASA$ computed using Rosetta InterfaceAnalyzer (see paragraph below). |

For each of the described features, we also computed the corresponding “normalized” feature, obtained by subtracting the average of the same feature across the 13 structures involving the same GPCR.

### Model training and testing

We used the state-of-the-art tabular foundation model Tabular Prior-data Fitted Network (TabPFN) through the tabpfn python package (version 2.0.9)^34^. The model used as input the features described in the previous section and the training objective was **max(log_10_(*c*),0.1)** where *c* is the GPCR-G protein coupling activity measured as percent of the one of the primary transducers (as provided by GproteinDb^17^).

We employed a repeated 5-fold cross-validation to ensure a robust evaluation of the model’s predictive performance. The data were partitioned at the G-protein-coupled receptor (GPCR) level. Specifically, we divided the 213 GPCRs in the training set into five distinct groups. For each fold, the model was trained on data related to the GPCRs in four of the groups and then tested on the data from the remaining group. To ensure the stability of the results, this entire 5-fold cross-validation process was conducted five times on independent training/test splits.

To determine the optimal cutoff for distinguishing positive from negative binders, we evaluated the model on the validation set. Model predictions were exponentiated to return to the original percentage scale. We then computed the Matthews Correlation Coefficient (MCC) using integer thresholds from 1 to 20 (Supplementary Figure 2A) and identified 4 (representing 4% of primary transducer activity) as the threshold that maximized the MCC.

### PRECOGx predictor model retraining

PRECOGx was retrained and tested following the same protocol as in the original paper^19^ with the same GPCR sequence representation through the protein language model ESM1b, but using the same GPCRdb dataset for training as in the new predictor.

### Feature importance computation

SHAP (SHapley Additive exPlanations)^35^ values were computed using the shap Python package (version 0.48.0) through the tabpfn-extension Python package (version 0.1.0). It employs the PermutationExplainer to approximate feature contributions for the TabPFN model. SHAP values were calculated for the held-out samples in each fold of the leave-one-G-protein-out cross-validation and for the IUPHAR test set and subsequently concatenated to form a single dataset for analysis.

### GPCRome-wide prediction of G protein complexes

The model was fitted on the entire dataset of experimentally tested GPCR-G protein pairs from GproteinDb (1,714 datapoints) and subsequently used to predict the coupling for the remaining 3,395 non-olfactory GPCR-G protein pairs and the 5,304 olfactory GPCR-G protein pairs. The threshold used to discriminate between positive and negative pairs was 4, chosen to maximize the accuracy of the predictions (see paragraph “Model training and testing”).

### Structure-based phylogenetic analysis

We retrieved 796 predicted structures of the receptors considered in this study from AlphaFold DB (v4)^74^. The 5 receptors exceeding 2700 amino acids (ADGRG4, ADGRV1, CELSR1, CELSR2, CELSR3) were excluded due to the unavailability of full-length predictions. Structure-based alignments were generated using the easy-msa module of FoldMason^37^ (Release 3-954d202) with default parameters, while sequence-based alignments were performed using Clustal Omega^75^ (5 iterations, --full option). All alignments were trimmed using trimAl (v1.5.rev0)^76^ with a gap threshold of 35% as in ^37^.

Phylogenetic trees were constructed using IQ-Tree v2.0.3^77^ with 1000 ultrafast bootstrap replicates^78^. The best substitution model was determined using ModelFinder^79^. For the structure-based 3Di alignment, we fixed the substitution matrix to that of ^80^, optimizing only character frequencies and rate heterogeneity (best model: 3Di+F+R10). For amino acid alignments, ModelFinder selected JTT+F+R10 as the best model for both the structure-based (FoldMason) and sequence-based (Clustal) MSAs.

The retention index was calculated using an in-house script that used the Fitch algorithm^81^ to compute the parsimony score on a given tree, using GPCR families or classes as characters. The retention index is then computed as defined in ^38^.

To compute the correlation between phylogenetics and coupling preferences, we defined for each receptor a coupling vector, that is, the 13-dimensional vector of coupling (experimental if available, predicted otherwise) to the G proteins considered in this work. The correlation between phylogenetics and coupling preferences was computed as the Pearson correlation coefficient between the phylogenetic distance matrix and the matrix of the Euclidean distances between the log-scaled coupling vectors.

### TGFα shedding assay

TGFα shedding assay was performed as described previously^8^ with minor modifications. Gαq/11/12/13-deficient HEK293 cells were seeded in a 6-well culture plate (Greiner Bio-One) at a concentration of 2.5 × 105 cells/mL (2 mL per well hereafter) in DMEM (Nissui Pharmaceutical) supplemented with 5% (v/v) FBS, glutamine, penicillin, and streptomycin, one day before transfection. The transfection solution was prepared by combining 5 µL of 1 mg/ml polyethylenimine Max solution (Polysciences) and a plasmid mixture consisting of 500 ng alkaline phosphatase (AP)-tagged transforming growth factor-α (AP-TGFα; human codon-optimized), 200 ng QRFPR plasmid and 10 ng chimeric Gαq construct with a substitution of the last 6 amino acids in 200 µL of Opti-MEM I Reduced Serum Medium (Thermo Fisher Scientific). One day after incubation, the transfected cells were harvested by trypsinization, neutralized with DMEM containing 5% (v/v) FCS and penicillin–streptomycin, washed once with HBSS containing 5 mM HEPES (pH 7.4), and resuspended in 6 ml of the HEPES-containing HBSS. The cell suspension was seeded into a 96-well plate at a volume of 80 µL (per well hereafter) and incubated for 30 minutes in a CO2 incubator. QRFP26 was serially diluted in assay by 3.2-fold (final concentrations ranging from 0.32 nM to 3.2 µM) and added to the plate with 10 µL per well. After 1 h incubation, the cell plate was centrifuged and conditioned media (80 µL) was transferred to an empty 96-well plate. AP reaction solution (10 mM p-nitrophenylphosphate (p-NPP), 120 mM Tris–HCl (pH 9.5), 40 mM NaCl, 10 mM MgCl2) was dispensed into the cell culture plates and plates containing conditioned media (80 µL). Absorbance at 405 nm was measured before and after a 2 h incubation at room temperature using a microplate reader (SpectraMax 340 PC384; Molecular Devices). The ligand-induced AP-TGFα release signals were fitted to a four-parameter sigmoidal concentration-response curve (Prism 10 software, GraphPad Prism) and logarithm of relative Emax/EC50 (Log RAi) was calculated as described in previously^8^.

### DNA plasmids, cell cultures, and transfections

HEK293T/17 cell line was purchased from ATCC (CRL-11268) and maintained in complete medium containing DMEM (Gibco, #11965118), 10% FBS (Biowest, #S1520), 1% non-essential amino acids (Gibco, #11140-050), 1% GlutaMAX (Gibco, #35050061), 1% sodium pyruvate (Gibco, #11360070), penicillin 100 units/ml and streptomycin 100 mg/ml (Gibco, 15140-122), and amphotericin B 250 g/ml (ThermoFisher, 15290-018) at 37°C and 5% CO_2_. Cells were routinely monitored for mycoplasma contamination. Two million cells were seeded in each well of 6-well plates in medium without antibiotics for 4 hours and then transfected with a 1:0.5 ratio of DNA plasmid (2.5 mg) and polyethylenimine (1.25 mL at 1 mg/mL) (Polysciences, #23966-100). FLAG-GPR3, FLAG-GPR6, FLAG-GPR12, FLAG-GPR50, FLAG-GPR55, FLAG-GPR37, FLAG-GPR37L1, and FLAG-GPR132 were obtained by removing the V2-tail, TEV, and tTA sequences from corresponding plasmids in the PRESTO-Tango library (Kit #1000000068). GRM2 and LPAR2 were a generous gift from Dr. Kirill Martemyanov (University of Miami, Florida). Plasmids encoding 5HT2A and 5HT4B were obtained from cDNA resources (www.cdna.org). The various reporters used, CRE-Nluc, NFAT-Nluc, and SRE-Fluc luciferase, were previously described in ^82,83^. The plasmid encoding GsGo was purchased from Addgene (#109375). For CRE-Nluc and NFAT-Nluc reporter assays, a firefly luciferase under the control of the constitutively active promoter thymidine kinase was used as a normalizer (pFL-tk). Similarly, pRL-tk encoding a Renilla luciferase was used as a normalizer for the SRF-RE reporter. Sequences of each construct were validated by whole plasmid sequencing at Eurofins Genomics.

### Luciferase Reporter Assays

G protein coupling for the orphan GPCRs tested was measured using four different luciferase reporters: CRE reporter for G_s_-mediated signaling, CRE reporter co-transfected with GsGo for G_i/o/z_-mediated signaling, NFAT reporter for G_q_-mediated signaling, and SRF-RE reporter for G_12/13_-mediated signaling^83^. For CRE reporter assays, the following plasmid ratios were used: 208 ng CRE-Nluc, 104 ng pFl-tk, and 1,772 ng receptor. 26 ng of GsGo were co-transfected exclusively when testing for G_i/o/z_-mediated signaling. For the NFAT reporter assay, plasmids were transfected according to the following ratios: 138 ng NFAT-Nluc, 70 ng pFl-tk, and 2,010 ng of receptor. For the SRF-RE reporter, the following plasmid ratios were employed: 83 ng of SRF-RE, 83 ng of pRL-tk, and 2,330 ng of receptor. pcDNA3.1 was used to normalize total DNA amounts. The day after transfection, cells were serum-starved for 4 hours in 1 mL OptiMEM (Gibco) and harvested in BRET buffer (PBS supplemented with 0.5 mM MgCl_2_ and 0.1% glucose), centrifuged at 500 x *g* for 5 minutes, and resuspended in 150 μL of BRET buffer. 30 μL of cells were plated in white 96-well plates (GreinerBio). 30 μL of Hikarazine™ Z103 (Synthelis #SYN90006) diluted 1:250 in BRET buffer were applied to detect NanoLuc emission, while 30 μL of BrightGlo (Promega) were used for firefly luciferase. For measuring pRL-tk, 10 µM coelenterazine-e (NanoLight Technology #355) prepared in BRET buffer was added as the substrate. Luminescence was acquired using a POLARstar microplate reader (BMG Labtech) until plateau values were reached. For each condition, the signal from the inducible reporter (CRE, NFAT, or SRF) was first normalized to the signal from the corresponding constitutively expressed luciferase (pFL-tk or pRL-tk) and subsequently normalized to pcDNA3.1. All measurements were performed at room temperature.

### Statistical analysis for luciferase reporters

Shapiro-Wilk normality test was used to assess the data gaussian distribution. Kruskal-Wallis vs pcDNA3.1 with Dunn’s multiple comparison was used for CRE, NFAT and CRE+GsGo reporters, while one-way ANOVA vs pcDNA3.1 with Dunnett’s multiple comparison was used for SRF reporter. Positive controls for reporters (5-HT4B for CRE, GRM2 for CRE + GsGo, 5-HT2A for NFAT, LPAR2 for SRF-RE) were excluded from the analysis

### Co-expression analysis

We obtained bulk RNA-seq data from the TCGA–TARGET–GTEx dataset available through the UCSC Xena Browser^84^. All pediatric samples (i.e., data from TARGET) were excluded. For tumor samples, only primary tumor samples were retained. After filtering, the total number of samples was 16,593 (9,181 tumor and 7,412 normal).

We identified and removed outlier samples for which any of the following metrics fell outside the specified ranges: number of detected genes (minimum 20,000; maximum 40,000), total read counts (minimum 10 million; maximum 120 million), and proportion of counts attributable to the 100 most highly expressed genes (no minimum; maximum 90%). These thresholds were determined by visual inspection of the corresponding distributions.

Next, the number of detected genes and the total number of counts were log1p-transformed, and adaptive filtering was applied independently to each tissue by removing samples that were more than five median absolute deviations (MADs) from the tissue-specific median for any of the metrics. Within each tissue, genes with fewer than 15 counts in at least 75% of samples were considered lowly expressed and excluded from downstream analyses. The data were normalized using the DeSeq2 variance stabilizing transformation^85^ implemented in PyDESeq2 (v0.4.4)^86^. Subsequently, principal component analysis was performed using the first 10 principal components, and samples whose Euclidean distance from the tissue-specific centroid exceeded five standard deviations were removed. Finally, Pearson correlation matrices were computed for each tissue using the PyWGCNA Python module^87^.

### Coupling map analysis of the co-expression network

We integrated GPCR co-expression results from GTEx (healthy tissues) and TCGA (tumor types) with a combined GPCR–G protein coupling resource (predicted and literature-curated). For all downstream analyses, we retained only edges with positive co-expression (co-expression > 0) and strong coupling support (coupling score ≥ 4). To enable cross-cohort comparisons, we harmonized sample categories by mapping each GTEx tissue to its corresponding TCGA cancer type (matched GTEx–TCGA categories).

For GTEx, we quantified GPCR family-level patterns of co-expressed and coupled receptors across tissues. Specifically, we summarized (i) global distributions of coupled/co-expressed GPCR classes across all tissues, and (ii) within-tissue family composition, computed as the proportion of GPCRs per family among all coupled/co-expressed GPCRs in that tissue. Family-level analyses were restricted to GPCR families with at least four members to reduce noise from sparsely represented families. We additionally summarized how widely each family appears across tissues by counting the number of tissues in which the family was detected, and highlighted families with the most restricted tissue distribution.

To compare GTEx and TCGA directly, we performed matched-category analyses of dataset content at multiple resolutions. We quantified differences in (i) the number of unique coupled GPCRs and (ii) the number of unique coupled GPCR–G protein coupling pairs (edges) observed in each dataset.

We assessed differences in dataset coverage between GTEx and TCGA using a presence/absence definition derived from family-level edge counts within matched categories. A GPCR family was considered present in a matched category if it had at least one GPCR–G protein edge (GTEx_n_pairs > 0 for GTEx; TCGA_n_pairs > 0 for TCGA). For each family, coverage was defined as the number of matched categories in which that family was present, computed separately for GTEx (***n_GTEX_***) and TCGA (***n_TCGA_***). Differences in coverage were summarized using a symmetric coverage metric: **Δ*cov_sym_*= (*n_TCGA_* − *n_GTEX_*)/(*n_TCGA_*+*n_GTEX_*)**. Positive values indicate greater coverage in TCGA, negative values indicate greater coverage in GTEx, and values near zero indicate similar coverage.

We defined the G protein coupling repertoire as the combinations of coupled G protein families, found co-expressed in each GTEx and TCGA tissues with cognate receptors. We finally decomposed these differences by counting dataset-specific GPCR–G protein edges and by summarizing global coupling shifts at the G protein family level using a Sankey diagram to visualize transitions between GTEx and TCGA.

### Software

We employed ChimeraX^88^ (v1.9) to generate all 3D cartoon representations with the exception of Figure 2F that was generated using Pymol (v2.5.0). We employed customized scripts in python (version 3.11.11), using matplotlib (v3.10.0), seaborn (v0.13.2), and biopython (v1.85) libraries. We calculated residue-residue contacts by using a customized script derived from the CIFPARSE-OBJ C + + library (https://mmcif.wwpdb.org/docs/sw-examples/cpp/html/index.html).

## Acknowledgments

F.R. was supported by the Italian Ministry of University and Research through the Department of Excellence “Faculty of Sciences” of Scuola Normale Superiore. The research leading to these results also received funding from the Italian Association for Cancer Research (AIRC) under My First AIRC Grant (MFAG) 2020 - ID. 24317 project. F.R. was also supported by the Project granted by Next Generation EU – National Recovery and Resilience Plan (Piano Nazionale di Ripresa e Resilienza, NRRP) – Mission 4 Component 2 Investment 1.4 – Ministry of University and Research (MUR) Call N. 3277 Project Code ECS_00000017 MUR Directoral Decree n.1055, 23 June 2022, CUP B83C22003930001, project title “Tuscany Health Ecosystem – THE”, Spoke 8. F.R. also received funding from the ImmunoHub project (“Immunoterapia: cura e prevenzione di malattie infettive e tumorali (Immuno-HUB)”). We gratefully acknowledge the CINECA award, in collaboration with AIRC, for the availability of high-performance computing resources and generous support. We gratefully acknowledge the computational resources of the Center for High-Performance Computing (CHPC) at Scuola Normale Superiore. A.I. was funded by KAKENHI JP24K21281 and JP25H01016 from the Japan Society for the Promotion of Science (JSPS); JP22ama121038 and JP22zf0127007 from the Japan Agency for Medical Research and Development (AMED); JPMJFR215T and JPMJMS2023 from the Japan Science and Technology Agency (JST); The Uehara Memorial Foundation. CO was supported by NIH grant DC022104.

## Data and Code availability

Dataset are available at the following link: https://precogx.bioinfolab.sns.it/precog3D

The code of the model is available here: https://github.com/raimondilab/Precog3D

